# New lessons on TDP-43 from the killifish *N. furzeri*

**DOI:** 10.1101/2021.02.04.429704

**Authors:** Alexandra Louka, Sara Bagnoli, Jakob Rupert, Benjamin Esapa, Gian Gaetano Tartaglia, Alessandro Cellerino, Annalisa Pastore, Eva Terzibasi Tozzini

## Abstract

Frontotemporal dementia and amyotrophic lateral sclerosis are fatal and incurable neurodegenerative diseases linked to the pathological aggregation of the TDP-43 protein. This is an essential DNA/RNA binding protein involved in transcription regulation, pre-RNA processing and RNA transport. Having suitable animal models to study the mechanisms of TDP-43 aggregation is crucial to develop treatments against disease. We have previously demonstrated that the killifish *Nothobranchius furzeri* offers the unique advantage as a model system that it is an organism with compressed lifespan and a conserved ageing phenotype that develops within months, making this organism the available shortest-lived vertebrate with a clear ageing phenotype. Here, we show that the two paralogs of TDP-43 from the killifish *N. furzeri* share high sequence homology with the human protein and recapitulate its cellular and biophysical behaviour. We prove that, during ageing, *N. furzeri* TDP-43 spontaneously forms insoluble intracellular TDP-43 aggregates that have amyloid characteristics and colocalize with the stress granule core protein G3BP. Our results propose this organism as a valuable model of TDP-43-related pathologies and show that even minute differences between the human and *N. furzeri* proteins may help to shed new light onto the role of TDP-43 in RNA recognition and granule formation.

## Introduction

TAR DNA-binding protein 43 kDa (TDP-43) is an RNA-binding protein involved in RNA metabolism (Buratti and Baralle, 2001; Kim et al., 2010; Sephton et al., 2010; Ayala et al., 2011; Tollervey et al., 2011; Alami et al., 2014; Ishiguro et al., 2016). It was first described in 1995 as a protein associated with HIV transcription (Ou et al., 1995) and reconsidered later on as an important component of splicing (Buratti and Baralle, 2001; Polymenidou et al., 2011) and mRNA transport and translation (Ishiguro et al., 2016; Neelagandan et al., 2018; Chu et al., 2019). More recently, TDP-43 was involved in the relentless motor neuron disease amyotrophic lateral sclerosis (ALS) and in the distinct, but genetically linked, frontotemporal dementia (FTD) (Neumann et al., 2006; Mackenzie and Rademakers, 2008; Ayala et al., 2011). Hallmarks of these diseases are the presence of aberrant, polyubiquitinated and hyperphosphorylated cytosolic aggregates of TDP-43 in different areas of the central nervous system (Neumann et al., 2006; Prasad et al., 2019). TDP-43 aggregates are found in 97% of sporadic ALS and 45% of specific FTD cases. Increasing consensus is that TDP-43 aggregation is not coincidental, but represents the pathological mechanism of these diseases (Hergesheimer et al., 2019).

Stress granule formation is one of the many cellular protective mechanisms as a response to cellular stress (Anderson and Kedersha, 2009). Their formation is initiated by the oligomerization of the core proteins Ras GTPase-activating protein-binding protein 1 (G3BP) and Cytotoxic Granule Associated RNA Binding Protein (TIA1) whose expression is regulated by TDP-43 (Tourrière et al., 2003; Gilks et al., 2004; McDonald et al., 2011). TDP-43 can be found in stress granules under cellular stress conditions and TDP-43 positive pathological inclusions in post-mortem tissues from ALS patients are positive for stress granule markers (Liu-Yesucevitz et al., 2010; Parker et al., 2012).

TDP-43 is a modular protein that binds UG-rich RNA sequences. It is composed of a partially unfolded N-terminal domain linked to two RNA-binding RRM repeats (RRM1 and RRM2) and followed by a C-terminal tail. Until recently, this region was thought to be the hotspot for protein aggregation, because it hosts most of the pathologic mutations (Berning and Walker, 2019) and contains a prion-like sequence (Louka et al., 2020). More recently, we demonstrated that other regions, including the RRM motifs, can play an important role in aggregation and pathology (Zacco et al., 2018). This evidence agrees with a role of RNA in the aggregation properties of this protein (Gotor et al., 2020; Loganathan et al., 2020). We also demonstrated that binding to short RNA aptamers can efficiently prevent TDP-43 aggregation (Zacco et al., 2019).

Several animal models have been developed to permit the study of TDP-43-mediated diseases, starting from C. *elegans* to mouse (Liu et al., 2013). One of the limitations that these systems offer is that either they are invertebrates or, if vertebrates, their relatively slow ageing dictates the pace by which studies can progress. Recently, a new vertebrate model organism was introduced that is based on the killifishes of the genus *Nothobranchius*, colloquially called annual fishes, and in particular the species *N. furzeri* (Cellerino et al., 2016). Killifishes are native to African savannah. Due to the environmental constraints in which this organism evolved to follow the periods of water abundance and depletion, *N. furzeri* matures within two weeks (Vrtílek et al., 2018), after which mortality increases rapidly. Also in captivity, lifespan is limited to 3-7 months depending on the genotype (Valdesalici and Cellerino, 2003; Terzibasi et al., 2008). The complete genome of *N. furzeri* is available and annotated (Reichwald et al., 2015; Valenzano et al., 2015). Techniques for reverse genetics are available (Harel et al., 2015) making the killifish a system potentially suitable for elucidating the pathologic mechanisms of disease and for screening new compounds. It was indeed demonstrated that the remarkably short lifespan of *N. furzeri*, currently the shortest-life vertebrate model organism available, recapitulates the main hallmarks of vertebrate ageing (Harel et al., 2015; Cellerino et al., 2016). *N. furzeri* was demonstrated to be an invaluable tool for studies in disparate branches of investigation such as evolutionary genomics (Reichwald et al., 2015; Valenzano et al., 2015; Sahm et al., 2017; Cui et al., 2020), regenerative medicine (Wendler et al., 2015), developmental biology (Dolfi et al., 2019; Hu et al., 2020), pharmacology (Valenzano et al., 2006; Baumgart et al., 2016) and ecotoxicology (Philippe et al., 2018). It was also shown that *N. furzeri* presents spontaneous age-dependent gliosis (Tozzini et al., 2012), neuronal protein aggregation and loss of stoichiometry of protein complexes in the brain that is triggered by early impairment of proteasome activity (Kelmer Sacramento et al., 2020) with selective age-dependent degeneration of dopaminergic and noradrenergic neurons in the midbrain (Matsui et al., 2019). It was showed that the fish presents inclusion bodies containing α-synuclein as those found in Parkinson patients and age-dependent degeneration of dopaminergic and noradrenergic neurons (Matsui et al., 2019). Genetic depletion of α-synuclein ameliorated symptoms demonstrating a causal link between aggregation and neurodegeneration.

In the present study, we explored the suitability of *N. furzeri* for the study of TDP-43-related pathologies. *N. furzeri* contains two TDP-43 paralogs, which we call hereafter Nfu_TDP-43 and Nfu_TDP-43L for *N. furzeri* TDP-43 and TDP-43-like proteins. We used a complementary approach based on *in silico, in vitro* and *ex vivo* techniques that allowed us to validate our results both at the cellular and at the protein level. We compared *in silico* and *in vitro* the tendency to bind RNA and aggregate of the *N. furzeri* proteins and their isolated domains with those of human TDP-43 (Hsa_TDP-43). We then demonstrated that the *N. furzeri* TDP-43 paralogs are able to aggregate spontaneously in the intact animal producing inclusions that are similar to those observed in humans. The process is ageing-related as it was detected only in old animals. Finally, we showed that the *N. furzeri* TDP-43 proteins co-localize in the fish with the stress granule-contained G3BP, indicating that also *N. furzeri* TDP-43 is able to segregate in stress granules.

Taken together, our data show that *N. furzeri* is a unique model system able to recapitulate most of both the *in vitro* and cellular phenotype observed in TDP-43 human diseases. This evidence opens the possibility to use *N. furzeri* as a powerful model in ALS studies and provides new information on the RNA-to-TDP-43 functional relationship.

## Materials and Methods

### Recombinant protein production

Recombinant constructs encompassing the RRM1-2 and C-terminal domains of Hsa_TDP-43 (K102–Q269, Q269-H414 and A315–H414, Uniprot entry: Q13148), Nfu_TDP-43 (K103– P269 and P269–M403, NCBI Ref: XP_015823124.1), and Nfu_TDP-43L (K103–Q269 and Q269–M405, NCBI Ref: XP_015814443.1) were cloned in a pET-SUMO plasmid. The *N. furzeri* sequences were only predicted and could thus be splicing variants as we could not exclude that there might be more isoforms for each gene. The cDNA was purchased from GenScript with codon optimization for *E. coli* expression. The plasmids were prepared by the Gibson’s assembly strategy (Gibson et al., 2009). The constructs were co-expressed in pETSUMO plasmids with an N-terminal hexa-histidine tag, a SUMO tag with a modified polylinker site and a cleavable tobacco etch virus (TEV) protease site. The proteins were purified as previously described (Zacco et al., 2019). In short, the plasmids were transformed in NiCo21(DE3) cells which were grown overnight at 37 °C in Luria–Bertani (LB) medium containing 50 μg/ml kanamycin up to a 0.7 optical density recorded at 600 nm, before inducing expression by adding 0.5 mM IPTG, overnight at 18 °C. The cells were then collected after centrifugation at 4000 rcf and lysed by sonication. The soluble fusion proteins were purified by nickel affinity chromatography (Super Ni-NTA agarose resin, Generon) and eluted from the column with high-salt phosphate buffer (10 mM potassium phosphate buffer, 150 mM KCL, 2.5 μM TCEP, at pH 7.2 for RRM1-2 and pH 6.5 for the C-terminal constructs) with the addition of 250 mM imidazole. The tag was cleaved off by incubating the construct with TEV (at a 1:10 protein to protease molar ratio). All samples were further purified with a second nickel affinity chromatography. Nucleic acids bound non-specifically to the protein constructs were removed by performing heparin affinity chromatography (HiTrap Heparin HP; GE Healthcare), while being connected to an Äkta pure system. This enables simultaneous observation of the absorbance at 260 and 280nm. The protein was eluted with using a high salt gradient (1.5 M KCl in a 10mM potassium phosphate buffer. Pure RRM1-2 domains of TDP-43 were obtained after size-exclusion chromatography with a HiLoad 16/60 Superdex75 prep grade in low-salt phosphate buffer (10 mM potassium phosphate buffer, 15 mM KCl, 2.5 μM TCEP pH 7.2 for RRM1-2 and 6.5 for the C-terminal constructs), aliquoted, flash-frozen and stored at −20 °C. The purity of the proteins obtained was checked by SDS-PAGE gels.

### ThT and Proteostat aggregation Assays

Aggregation kinetics of the isolated proteins were determined by the Thioflavin T (ThT)-binding assay followed by a BMG FLUOstar Omega microplate reader in Greiner Bio-One CELLSTAR plates. When not carried out immediately after purification, protein aliquots were rapidly thawed from the freezer, spun down, and filtered with a 0.2 μm syringe filter before each assay to ensure removal of any contingent pre-formed oligomer. The resulting concentration was determined. The solutions were then diluted to reach final concentrations of 10 μM for the RRM1-2 domains and 6 μM for the C-termini. The concentration of added ThT was 20 μM. The plate was sealed with an optic seal to avoid evaporation. The experiments were carried out at 37°C by recording the fluorescence intensity of ThT as a function of time for 48 h using 430 nm for the excitation wavelength and 485 nm for the emission. The experiments in which the fusion protein was cleaved directly in the plate were carried out by adding TEV protease directly to the protein samples using 1:10 enzyme to protein molar ratios.

When the experiments were carried out in the presence of RNA, the fluorescent dye used was ProteoStat, which, as opposed to ThT, does not interact with RNA. The aptamer was added to the protein in an RNA to protein 2:1 molar ratio using 10 μM concentrations of RRM1-2. The excitation wavelength was set at 500 nm and the emission at 600 nm.

All the reported data derive from at least three repetitions and are presented as percentage values, where 100% is the highest fluorescence intensity value registered during each set of assays.

### Circular Dichroism Measurements

Circular dichroism spectra were recorded on 10 μM samples with a JASCO-1100 spectropolarimeter. The spectra were acquired in 1 mm path length quartz cuvettes under a constant N2 flush at 4.0 L/min. All CD datasets were an average of thirty scans. For the determination of the global secondary structure, the far-UV (190–260 nm) spectrum was recorded at 20°C in low-salt phosphate buffer. Samples were gradually heated to 90°C (1°C/min) for the determination of the melting temperature (Tm). The ellipticity variation as expressed by the intensity of the band at 208 nm was followed as a function of temperature. The data were plot and the Tm calculated according to literature (Greenfield, 2006). Control CD spectra were acquired after the temperature was brought back to the original 20°C (1°C/min). The spectra were corrected for the buffer signal and expressed as mean residue molar ellipticity θ (deg*cm^2^*dmol^-1^).

### Animal housing and tissue preparation

The animal facility was licensed by the Italian Ministry of Health (D.M. n. 80/2013 - A). All procedures for fish culture and organ harvesting were approved by the Animal Welfare Committee of the University of Pisa and notified in advance to the Ministry of Health in accordance with the European Directive 2010/63/EU and the Italian Law.

Brains from 5 and 27 weeks old animals were micro-dissected and processed for immunohistochemistry protocols: tissues were first fixed with paradormaldehyde (PFA) 4% in PBS (O/N at 4 °C), then cryo-protected with sequential immersions in sucrose 20% and 30% until the tissue precipitated to the tube bottom (a minimum of 6 hours/each steps). Finally, the tissues were singularly included in Tissue-Tek®O.C.T.™ Compound (Sakura) and 25 tick sections were cut on Superfrost Plus adhesion slides (Thermo Fisher Scientific).

### Immunofluorescence and Proteostat aggresome assay

We performed immunofluorescence experiments on cryo-sections of 25 microns and proceeded as previously described (Tozzini et al., 2012). Briefly, we washed the sections in PBS to remove the cryo-embedding medium. We then performed an acid antigen retrieval step (10 mM Trisodium citrate dehydrate, 0.05% tween, at pH 6) and (when required) stained the section with aggresome (ProteoStat Aggresome Detection Kit, Enzo Life Sciences Inc.) as previously described in details (Shen et al., 2011). We applied a solution 1:2000 of aggresome dye in PBS for 3 minutes, rinsed the samples with PBS and left the sections immersed in a solution of 1% acetic acid for 40 minutes. We applied blocking solution (5% BSA, 0.3% Triton-X in PBS) for 2 h. Primary antibodies at proper dilution were added in a solution of 1% BSA, 0,1% triton in PBS, and left overnight at 4°C (**Table 3**). The day after we applied secondary antibodies at a 1:400 dilution in the same solution. After 2h, slides were washed three times with PBS and mounted with a specific medium added with nuclear staining (Fluoroshield DAPI mounting medium, from Sigma-Aldrich).

### 3D visualization of doughnut-like cells in N. furzeri

We used the AbSca/e method to visualize the cells presenting an abnormal nuclear distribution of TDP-43 in *N. furzeri*. This is a clearing and staining approach for whole mount tissues that avoids tissue expansion and preserves lipids (Hama et al., 2015). It has superseded the original Sca/e method which was much slower. The technique was adapted to the considerably smaller size of *N. furzeri* brains (**Tables 1 and 2**). In short, the brains were fixed after dissection in 4% PFA overnight and adapted with a S0 solution for 18 h at 37°C. The samples were then permeabilized with sequential incubation in A2-B4(0)-A2 solutions at 37°C. After the permeabilization steps, samples were de-Scaled through incubation in PBS for 6 h at room temperature followed by incubation with TDP43 primary antibody in an AbSca/e solution for 3 days at 4°C, rinsed twice in AbSca/e for 2 h each and incubated with a secondary antibody for 18 h at 4°C. The samples were then rinsed for 6 h in AbSca/e and subsequently in the AbRinse solution twice for 2 h. After a re-fixation step in 4% PFA for 1 h and a rinse in PBS for an additional hour, samples were finally clarified in Sca/e S4 for 18 h at 37° and maintained in Sca/eS40 at 4° until imaging. The Sca/eS4 solution was also used as imaging medium.

**Table 1 –.** AbSca/e staining procedure. This is the procedure we adapted from Hama et al., 2015 for clearing and staining of *N. furzeri* brains. In the table are reported the solution required for each step with the time and temperature of incubation. ON: over night, RT: room temperature.

| Step | Solution | Timing (approx.) | Temp. |
| --- | --- | --- | --- |
| Fixation | 4% PFA | ON | 4°C |
| Adaptation | Sca/e S0 | 18h | 37° |
| Permeabilization | Sca/e A2 | 36h | 37° |
|  | Sca/e B4(0) | 24h | 37° |
|  | Sca/e A2 | 12h | 37° |
| Descaling | PBS | 6h | RT |
| Immunostaining | AbSca/e + primary antibody | 3 days | 4° |
|  | AbSca/e | 2h (2x) | RT |
|  | AbSca/e + secondary antibody | 18h | 4° |
| Wash | AbSca/e | 6h | RT |
| Rinse | AbRinse | 2h (2x) | RT |
| Refixation | 4% PFA | 1h | RT |
| Wash | PBS | 1-2h | RT |
| Clearing | Sca/e S4 | 18h | 37° |
| Mounting | Sca/e S4 |  | 4° |

**Table 2 –.** Composition of the solution used for the AbSca/e protocol. The composition is taken as indicated in Hama et al., 2015.

| Ingredients | Sca/eA2 | Sca/e B4(0) | Sca/eS0 | Sca/eS4 | AbSca/e | AbRinse |
| --- | --- | --- | --- | --- | --- | --- |
| D-(-)-sorbitol (w/v)% | - | - | 20 | 40 | - | - |
| Glycerol (w/v)% | 10 | - | 5 | 10 | - | - |
| Urea (M) | 4 | 8 | - | 4 | 0.33 | - |
| Triton-X-100 (w/v)% | 0.1 | - | - | 0.2 | 0.5 | 0.05 |
| Methyl-β-cyclodextrin (mM) | - | - | 1 | - | - | - |
| γ-Cyclodextrin (mM) | - | - | 1 | - | - | - |
| N-acetyl-L-hydroxyproline (w/v)% | - | - | 1 | - | - | - |
| DMSO (w/v)% | - | - | 3 | 25 | - | - |
| PBS | - | - | 1x | - | - | 0.1x |
| BSA (w/v)% | - | - | - | - | - | 2.5 |

**Table 3 –.** List of antibodies utilized in the immunofluorescence experiments.

| **Antibody** | **Product type** | **Working dilution** | **Company (Cat N.)** |
| --- | --- | --- | --- |
| TDP-43 | Rabbit Polyclonal | 1:1200 | Proteintech (10782-2-AP) |
| TDP-43 | Rabbit Monoclonal | 1:500 | AbCam (ab190963) |
| G3BP | Mouse Monoclonal | 1:500 | AbCam (Ab56574) |

### Computational tools

The catGRANULE software (http://service.tartaglialab.com/new_submission/catGRANULE) was used to predict the protein tendency to phase separate (Bolognesi et al., 2016). The results are available in https://tinyurl.com/y45hp82b. catRAPID *signature* was used to predict the propensity of TDP-43 to interact with RNA and identify RNA-binding domains (Livi et al., 2016). The software can be found at http://service.tartaglialab.com/new_submission/signature. The catRAPID *signature* results can be retrieved from https://tinyurl.com/y3ondta2, https://tinyurl.com/y3w57dsg and https://tinyurl.com/y5a6rb3a for Hsa_TDP4, Nfu_TDP-43L and Nfu_TDP-43 respectively. The catRAPID *omics* was employed to compute the binding targets in the human transcriptome (Agostini et al., 2013). The software can be found at http://s.tartaglialab.com/page/catrapid_omics_group. The catRAPID *omics* results are available at https://tinyurl.com/y2swjuxl for Hsa_TDP-43, Nfu_TDP-43L, Nfu_TDP-43 and *N. furzeri* methyltransferase (A0A1A8VE13 or TFB2M) The GU-rich sequences retrieved can be found at https://tinyurl.com/y5jd8wm5).

## Results

### *N. furzeri* and *H. Sapiens* TDP-43 have similar phase-separation propensities

Sequence analysis showed an impressive degree of sequence conservation between the *N. furzeri* and human TDP-43: Nfu_TDP-43 and Nfu_TDP-43L share 80% identity with each other, and 75% and 77% identity with Hsa_TDP-43 respectively (**Figure 1**). The homology is however not equally distributed. The N-terminus and the RRM domains are highly homologous, with hardly any indel (insertion and deletions) up to Q269. The only noticeable difference is an insertion of six residues in Nfu_TDP-43 near the interface between RRM1 and RRM2. More divergent are the C-termini with a degree of identity of 56% and 59% between the human protein and Nfu_TDP-43 and Nfu_TDP-43L respectively. The C-termini of the two *N. furzeri* paralogs share 63% identity.

**Figure 1 –.**
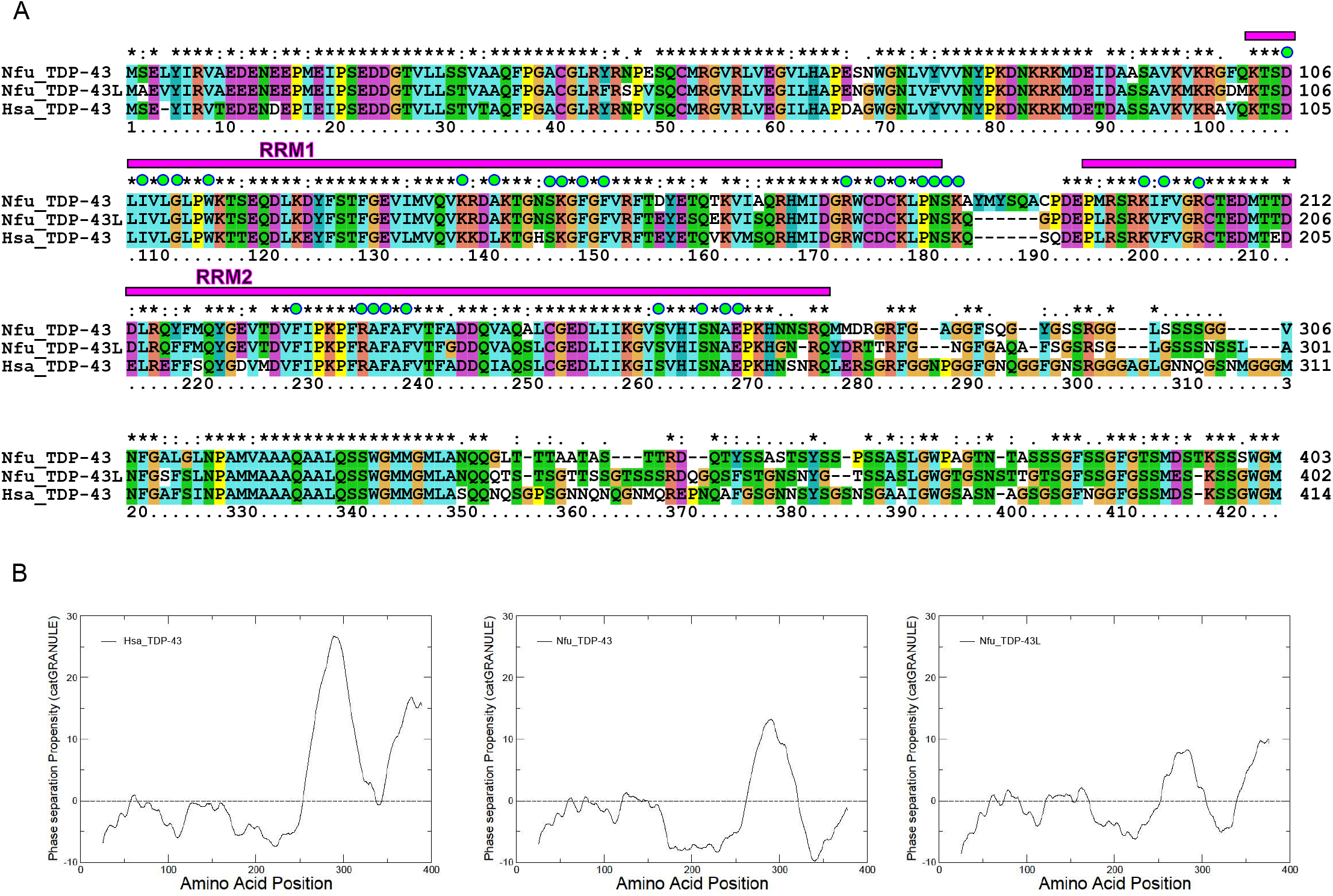
Bioinformatic comparison of *N. furzeri* and human TDP-43. **A)** Multiple alignment of the sequences produced by clustalx. The position of the two RRM domains is indicated with a magenta box. The amino acids in contact with the RNA aptamer observed in the solution structure (Lukavsky et al., 2013, 4bs2) are indicated with a green spot. **B)** catGRANULE profiles of the tendence of Hsa_TDP-43 (left), Nfu_TDP-43 (middle) and Nfu_TDP-43L (right) to have phase transitions.

We then reasoned that an important property of human TDP-43 is to phase separate and form stress granules. This property is thought to be mediated by TDP-43 C-terminus (Louka et al., 2020). Since this is the most divergent region amongst the three protein sequences, we compared the tendency *in silico* of the proteins to form granules by the catGRANULE approach (Bolognesi et al., 2016). This software takes into account structural disorder, nucleic acid binding propensity and amino acid patterns such as the presence of arginine-glycine and phenylalanine-glycine motifs to predict the tendency of a protein to coalesce in granules. We observed consistently a high tendency to phase separate in all three proteins which, as expected, is mostly localised in the C-termini with minor differences. The overall *cat*GRANULE scores of Hsa_TDP-43 and Nfu_TDP-43L are >1 indicating high-propensity to form liquid-like compartments, while Nfu_TDP-43 has a slightly smaller score of 0.8 (**Figure 1B**).

### The C-terminus of *N. furzeri* TDP-43 is highly aggregation-prone in *vitro*

We then characterized *in vitro* the isolated domains of the *N. furzeri* proteins using purified recombinant constructs with the aim of comparing their properties with those of Hsa_TDP-43 while being able to appreciate the individual contribution. We first analyzed the more distantly related C-terminal domains. The recombinant C-termini of Nfu_TDP-43 and Nfu_TDP-43L (starting from the end of RRM2) resulted soluble in bacteria and could be cleaved and purified from the SUMO tag although with tiny yields. Conversely, Hsa_TDP-43 could only be expressed in inclusion bodies as previously reported by independent studies (Li et al., 2018a; Shih et al., 2020). Rather than re-dissolving the protein from inclusion bodies, we decided to excise the glycine-rich region and produced a shorter fragment spanning residues 315-414 which resulted soluble. This construct did not contain the main hotspot for granule formation but incorporated many of the motifs thought to be crucial in protein self-association and phase separation (Mompeán et al., 2014, 2015; Li et al., 2018b, 2018a).

As in many traditional studies, we measured the kinetics of aggregation following the signal of a fluorescent dye (thioflavin T or ThT) detected at 485 nm. Since, however, all constructs resulted anyway little soluble demonstrating the strong intrinsic tendency of these regions toward aggregation, we carried out the assays in two different ways. We first used the fusion proteins without cleaving the constructs from the tag. Under the conditions of the assay, the fluorescent signal of all three proteins increased steeply within the first 2-3 h indicating fibre formation (**Figure 2**). After this time, the signal of Nfu_TDP-43L and Hsa_TDP-43 C-termini reached a plateau but not that of Nfu_TDP-43 whose signal decayed rapidly. This behaviour possibly indicated a stronger tendency of this protein fragment to precipitate. The Hsa_TDP-43 C-terminus reached comparatively higher fluorescence values and seemed to have a secondary increase after ca. 60 h. For comparison, we repeated the assay cleaving the proteins from the tag directly in the presence of ThT and followed their fluorescence signals versus time. We observed a qualitatively similar behaviour for the three constructs with a decrease of all signals. The signal of Nfu_TDP-43 C-terminus decreased appreciably faster.

**Figure 2 –.**
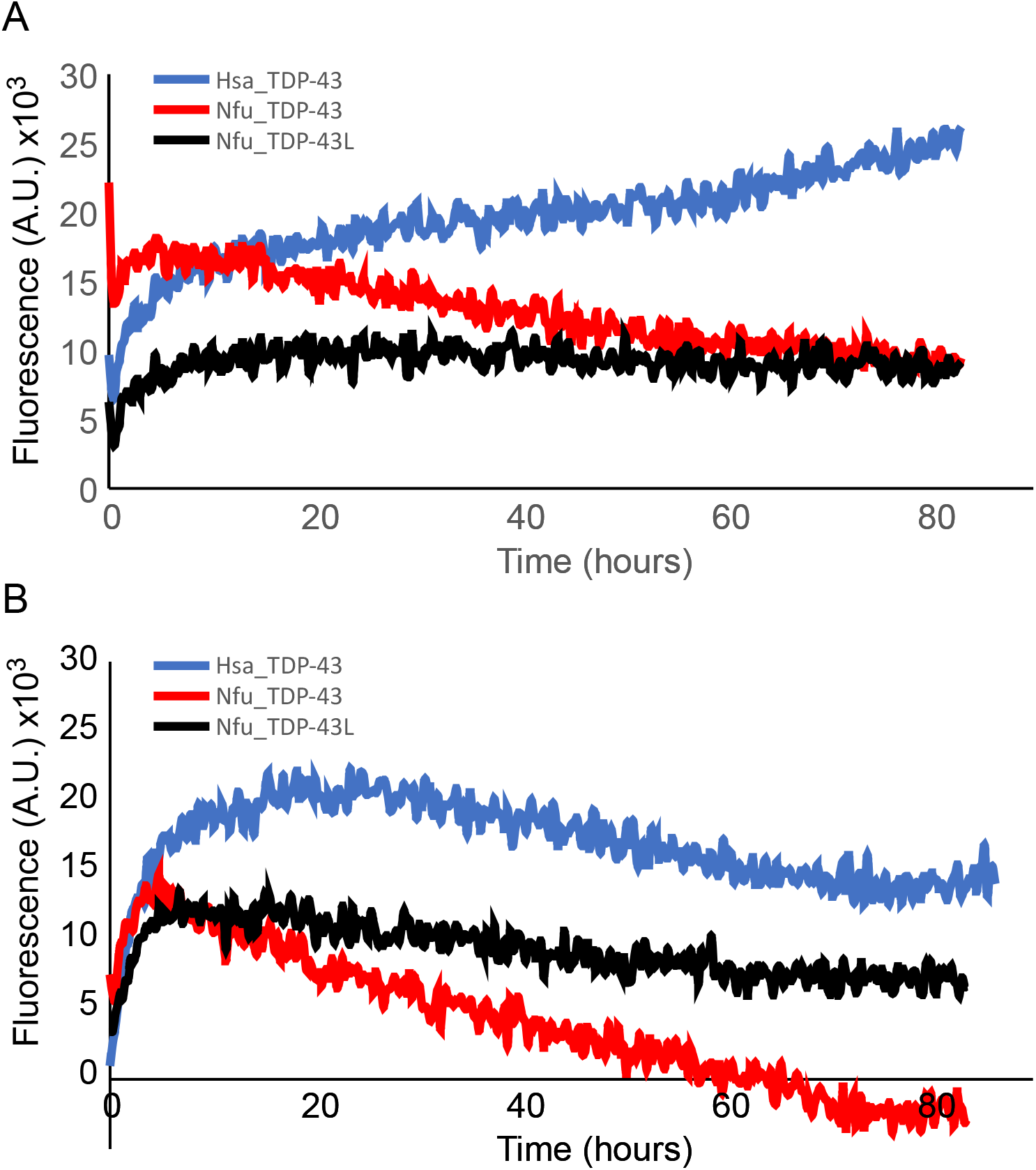
Comparison of the aggregation of the TDP-43 C-termini as detected by ThT aggregation assays. A) Aggregation assays carried out on constructs uncleaved from the SUMO tags. The protein concentration was 6 μM. B) The same as in A but cleaving the proteins -during the assay by TEV protease (1:10 protein to protease molar ratio). Drastic reduction of the signal of Nfu_TDP-43 indicates precipitation.

These data confirm an important role of the C-terminus in protein aggregation also for the *N. furzeri* proteins in full agreement with predictions. However, the possibility to produce the C-terminus in a soluble recombinant form using the *N. furzeri* sequences opens new avenues for the study of TDP-43 aggregation.

### The role of RRM1-2 in the aggregation of *N. furzeri* TDP-43

Similar experiments were carried out on constructs containing the two isolated tandem RRM domains (RRM1-2) to assess the contribution to aggregation of regions of the proteins other than the C-termini and, specifically, of the RNA-binding domains. Not unexpectedly, RRM1-2 proved to be more soluble than the C-terminus and could be efficiently produced in appreciable quantities and high purity. Overall, no major differences were observed amongst the constructs upon aggregation: In all three cases, the fluorescent signal increased as a function of protein concentration (**Figure 3A**), with aggregation that became progressively visible during the first 20 h of the assay. The curves of the two paralogs did not reach a clear plateau within the time of the experiment (3 days). Apart from these minimal differences, the curves exhibited similar shapes with the intensity of the human protein being noticeably higher than that of the *N. furzeri* constructs at comparable concentrations. No difference was observed when the assays were repeated in the presence of salt (15 mM KCl) (data not shown).

**Figure 3 –.**
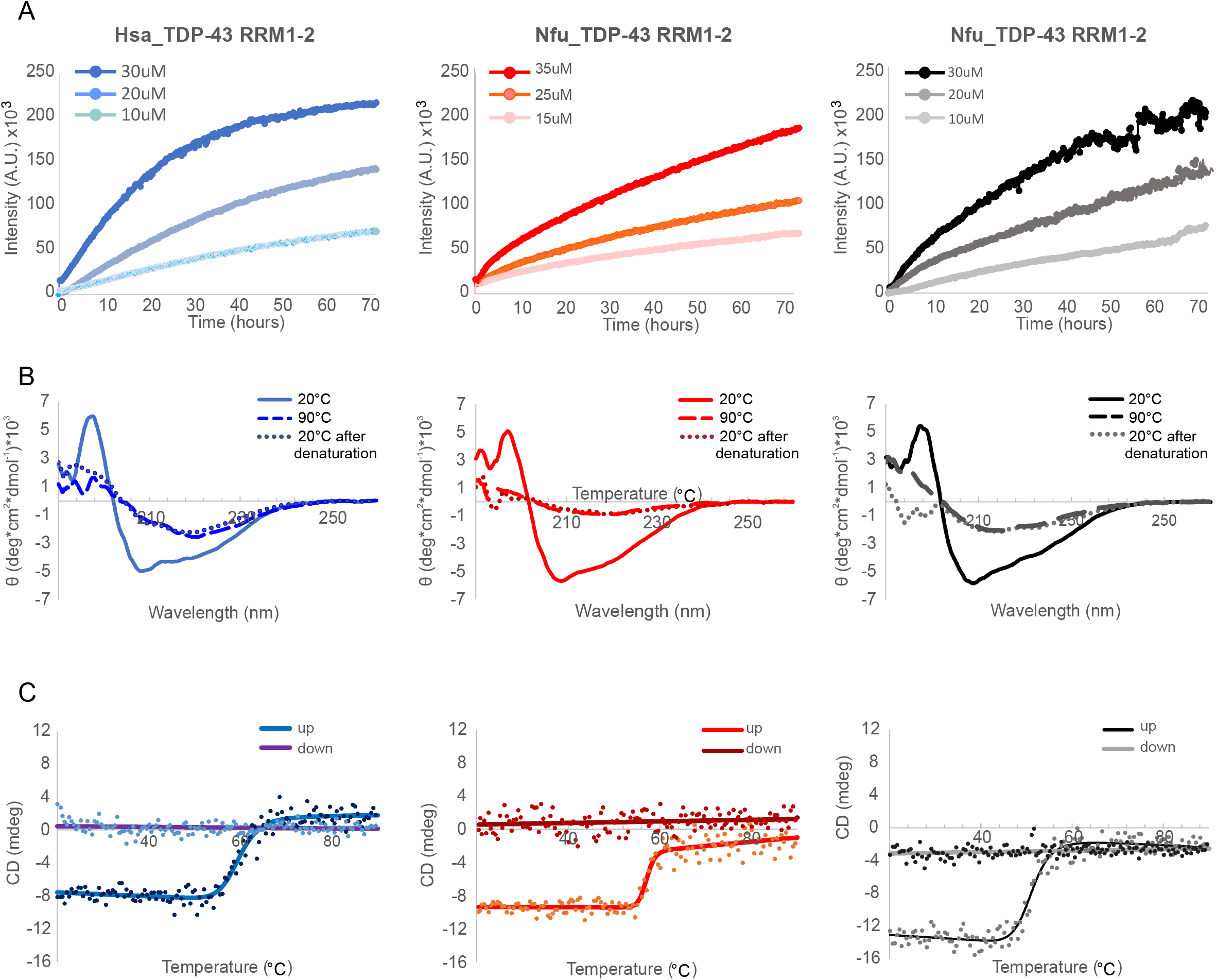
Comparison of the tendency to aggregate and adopt a β-rich structure of the RRM1-2 constructs. A) ThT aggregation assays at increasing protein concentrations. The concentration of ThT was 20 μM. B) CD spectra of the proteins (10 μM) at different temperatures. The spectra were corrected for buffer absorbance. C) Thermal stability of the three proteins under the same conditions as in B.

These results confirm an involvement in aggregation of the RNA-binding domains as already observed for the human protein (Zacco et al., 2018 and 2019) and demonstrate a qualitatively similar behaviour for the three proteins as expected from the high sequence homology of this region.

### *N. furzeri* TDP-43 RRM1-2 undergoes an irreversible conformational transition similar to the human protein upon thermal destabilization

To confirm the presence of a transition towards a β-rich structure upon protein destabilization which would testify misfolding, circular dichroism (CD) spectra of purified *N. furzeri* RRM1-2 (10 μM) were measured immediately after the final gel filtration step of purification (**Figure 3B**). The spectrum at 20 °C had two minima at 207 nm and 215 nm, a shape indicative of a mixed αβ conformation. The intensities correlated well with the known structure of this region (Lukavsky et al., 2013) and with previous studies (Zacco et al., 2018). When the temperature was gradually increased up to 90 °C, the two minima collapsed into a single shallow minimum at approximately 215 nm as expected for an increase in β-secondary structure. The spectra remained the same after returning at 20 °C indicating irreversibility of the process. We thus assumed that the change of the spectra must be due to a conformational transition which cooccurs with aggregation.

Thermal denaturation was then recorded following the intensity variations of the band at 222 nm as a function of temperature (**Figure 3C**). The curves indicated an irreversible transition with an apparent melting temperatures of 57.1 and 50.8 °C for Nfu_TDP-43 and Nfu_TDP-43L respectively to be compared with the melting of 59.6 °C for Hsa_TDP-43. The samples were then extracted from the cuvette and centrifuged at 16000 rcf. An absorbance spectrum between 600 nm and 200 nm of the soluble fraction showed no absorbance at 280 nm indicating almost complete precipitation of the proteins.

The three constructs have thus a similar behaviour also in terms of a α-to-β irreversible transition of the RRM domains triggered by heat destabilization although with minor differences in their resilience against aggregation.

### Similarities and differences of RNA-binding properties of *N. furzeri* and human TDP-43

We then compared *in silico* the RNA-binding propensities of the full-length proteins. We used catRAPID *signature* (Livi et al., 2016), a program based on physico-chemical and secondary structure properties and hydrophobicity profiles, to predict RNA-binding regions. catRAPID *omics* (Agostini et al., 2013) was used to identify RNA sequences able to bind. catRAPID *omics* ranks interactions according to a score (Bellucci et al., 2011) and estimates the binding potential through van der Waals, hydrogen bonding and secondary structure propensities of both protein and RNA sequences allowing identification of binding partners with 80% accuracy (Lang et al., 2019).

catRAPID *signature* predicted similar RNA-binding profiles along the protein sequences for the three proteins (**Figure 4A**). Two regions consistently exceeded the threshold that defines binding and they coincided, as expected, with the two RRM domains. Overall, however, the profiles of Nfu_TDP-43L and Hsa_TDP-43 were more conserved, especially over the RNA-binding domains. Other regions came out just around or above the threshold suggesting the presence of yet unidentified RNA-binding sites. We then ran *cat*RAPID *omics* against a human transcript library, imposing a high-confidence threshold (catRAPID scores > 2.5). We found 550 transcripts predicted to interact with the human protein. Nfu_TDP-43L was predicted to interact with 936 human transcripts (out of 10^5^ analysed) of which 550 were in common with Hsa_TDP-43. Nfu_TDP-43 was predicted to interact with the same 550 human RNAs but, in addition, with another set of 719 targets. In contrast, *N. furzeri* methyltransferase, used as a control, was predicted to interact with just 109 targets of which only 21 sequences being in common with Hsa_TDP-43.

**Figure 4 –.**
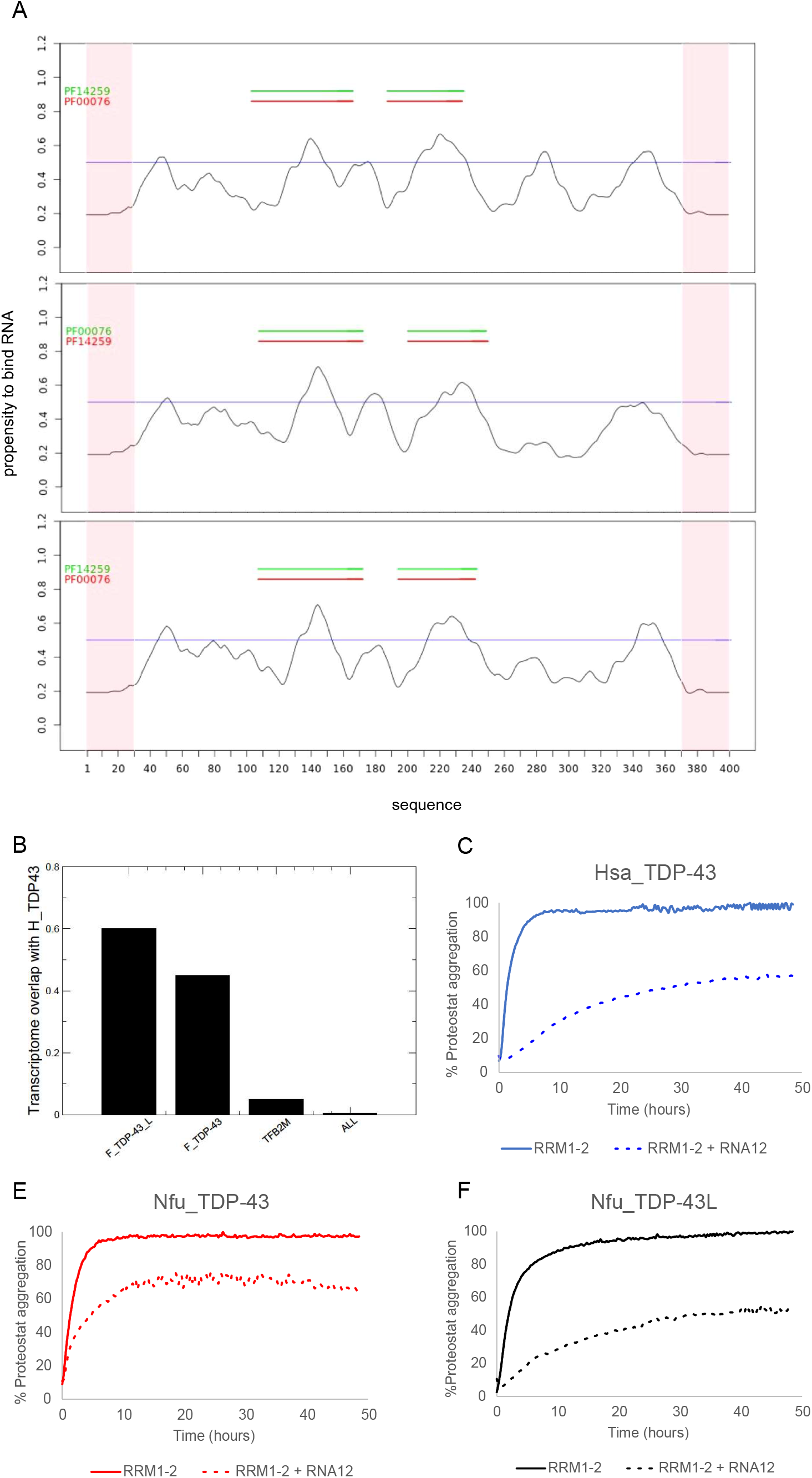
RNA-binding propensity of *N. furzeri* TDP-43 sequences as compared to Hsa_TDP-43. A) catRAPID *signature* prediction of the RNA-binding profile along the protein sequences of Hsa_TDP-43 (top), Nfu_TDP-43L (middle) and Nfu_TDP-43 (bottom). B) The Jaccard index indicates the predicted transcriptome overlap of other proteins with Hsa_TDP-43. The entire transcriptome ‘ALL’ contains 10^5^ transcripts. The trascripts of Nfu_TDP-43_L have the best overlap. Nfu_TDP-43 shows a slightly smaller coverage, while methyltransferase TFB2M, used as a control, has a negligible overlap. C,D,E) Comparison of the effects of the RNA12 aptamer on the tandem RRM1-2 domains of Hsa_TDP-43 (C), Nfu_TDP-43 (D) and Nfu_TDP-43L (E) on the aggregation properties of human and *N. furzeri* proteins followed by fluorescence using the Proteostat dye.

Overall, the Jaccard index, i.e. the ratio between the intersection and the union of two sets, considering enrichments of shared over total RNA interactions, is 550/(385+550) ≈ 0.60 for Nfu_TDP-43L, 550/(719+550) ≈ 0.45 for Nfu_TDP-43 and 21/(550+109) ≈ 0.05 for the control. This tells us that Hsa_TDP-43 and Nfu_TDP-43L have more similar propensities than Nfu_TDP-43 (**Figure 4B**). A Fisher exact test confirmed that the overlap between the *N. furzeri* and *H. sapiens* TDP-43 transcriptomes is significant (p-value < 0.00001; 10^5^ transcripts used as background). Analysis of the motifs present in the top 100 RNA targets shared between Nfu_TDP-43L and Hsa_TDP-43 indicated an enrichment of GU-rich sequences (CUGG[ACG][UA][CG], UG[CG]UG[ACG][UA] and CUGG[ACG]A) (Agostini et al., 2014) in agreement with what reported in the literature (Bhardwaj et al., 2013).

To have some validation of these results experimentally, we checked the effect of an RNA aptamer on the aggregation of the *N. furzeri* RRM1-2 constructs. We had previously demonstrated that an aptamer (RNA12), known to bind human RRM1-2 with nanomolar affinities (Lukavsky et al., 2013), is able to inhibit aggregation of this domain almost completely (Zacco et al., 2019). We thus probed the ability of this aptamer on the aggregation of the *N. furzeri* proteins following the fluorescence signal of the Proteostat dye. This fluorophore was chosen for these assays because ThT is known to bind to RNA resulting in interferences with the measurement (Zacco et al., 2019). Based on the high sequence homology and the specific conservation of the residues involved in interaction, we expected that *N. furzeri* TDP-43 proteins would have RNA-binding properties identical to the human protein. We found instead that RNA12 has similar inhibitory effects on Hsa_TDP-43 and Nfu_TDP-43L but a milder effect on Nfu_TDP-43, suggesting a lower affinity binding with the RNA aptamer (**Figure 4C**). This similarity between Hsa_TDP-43 and Nfu_TDP-43L binding ability is in agreement with our *cat*RAPID *omics* predictions.

These results give us some idea about the exquisite sensitivity by which even small differences in sequence can change the affinity for RNA of these proteins.

### TDP-43 progressively aggregates with ageing of *N. furzeri*

We then probed the localization of the protein distribution throughout the principal areas of the fish brain, such as telencephalon and optic tectum performing immunofluorescence experiments on 25 μm cryo-sections of *N. furzeri* MZM-04010 brains. We compared animals at 5 weeks, the age at which the animal reaches maturity, with animals at 27 weeks, when age-dependent mortality starts (Terzibasi et al., 2008). To verify the robustness of the staining pattern, we used two different antibodies raised against the human protein with different epitope specificities: a monoclonal (Abcam) and a polyclonal (Proteintech) rabbit antibody, respectively. The two different antibodies showed comparable staining patterns (**Figure 5**). The monoclonal antibody showed an overall cleaner signal, which was also more stable over prolonged storage. The staining associated with the polyclonal antibody was instead labile and decayed in a few days. For this reason, we decided to use the monoclonal antibody in all further experiments.

**Figure 5 –.**
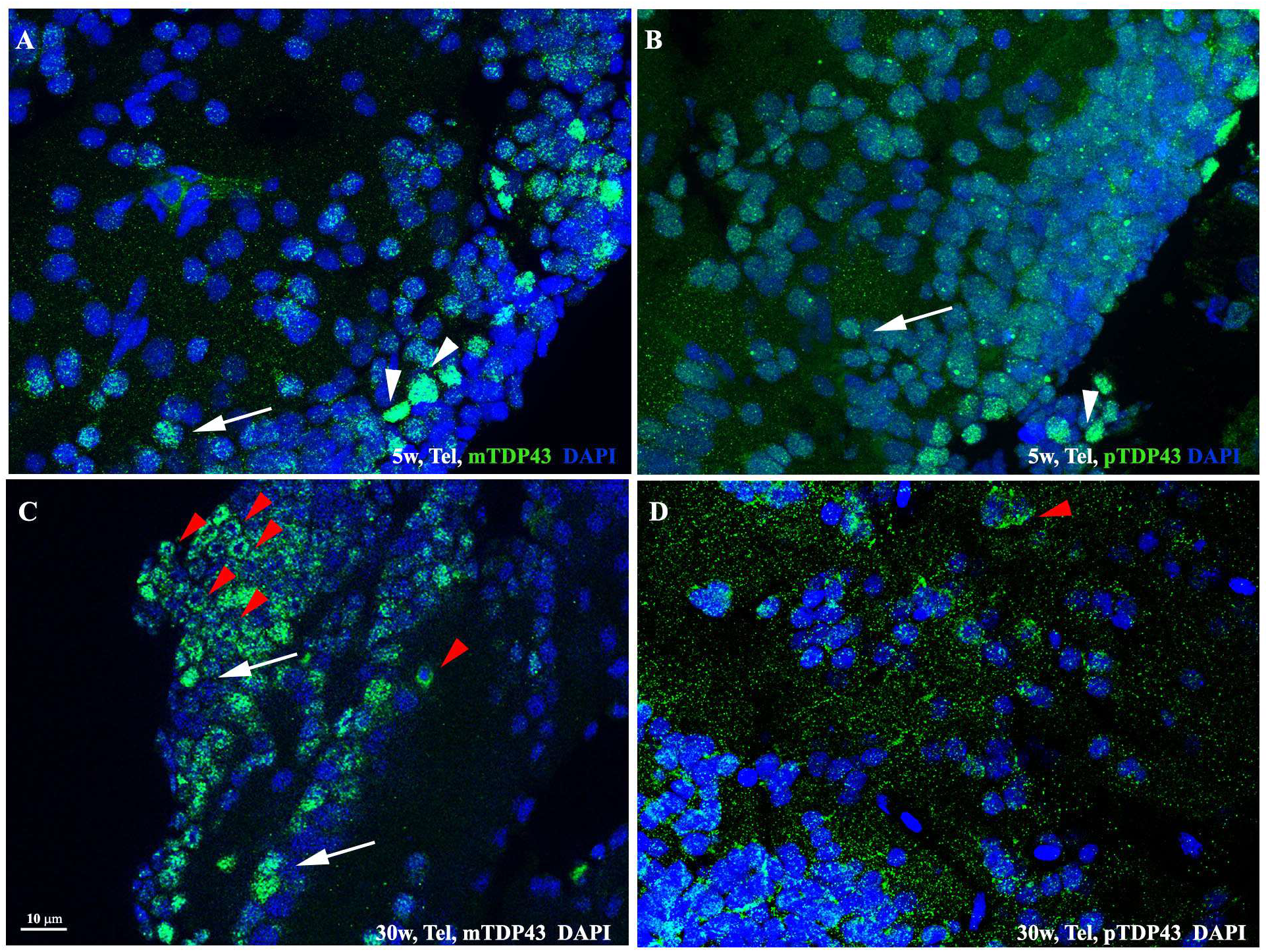
Immunostaining of TDP-43 with monoclonal (A,C) and polyclonal antibody (B,D). The staining resulted quite comparable in both cases with the presence of cells showing a more intense labelling (white arrowheads) and other with a less strong signal (white arrows). Both staining highlighted the presence of doughnut-like cells especially in samples of old fishes (panels C,D, red arrowheads). The scale bars are the same for all panels.

TDP-43 staining showed a variable level of diffuse expression in the cell nuclei of young animals (**Figures 5A, B**), presenting cells with brighter signal (white arrowheads), and others with a less intense TDP-43 positivity (white arrows). Staining in old tissues was characterized by the presence of cells with a peculiar endo-perinuclear concentration of protein labelling (**Figures 5C, D**, doughnut-like stained cells, red arrowheads). Doughnut-like stained cells were sporadically present in the young animals (data not shown). This fact, together with the presence of cells with diffuse labelling in the old tissue, suggested that the altered protein distribution has a progressive evolution over time.

To better appreciate the doughnut-like TDP-43 distribution in old animals, we performed whole mount brain staining (**Figure 6 and Movies S1 and S2**): we clarified and stained the whole *N. furzeri* brains, by using a Sca/e S immunofluorescence-optimized methodology (AbSca/e) (Hama et al., 2015), and registered 3D brain reconstructions by acquiring confocal serial stack images and processing them through the IMARIS Software. By comparison of the 3D representations of the tissues from young (**Figure 6Aa and Movie S1**) and old (**Figure 6Bb and movie S2**) animals, we observed an apparently higher proportion of doughnut-like cells (red arrowheads) in the telencephalic area of the old fish as compared to that of the young ones. Diffuse stained nuclei were detectable in both tissues.

**Figure 6 –.**
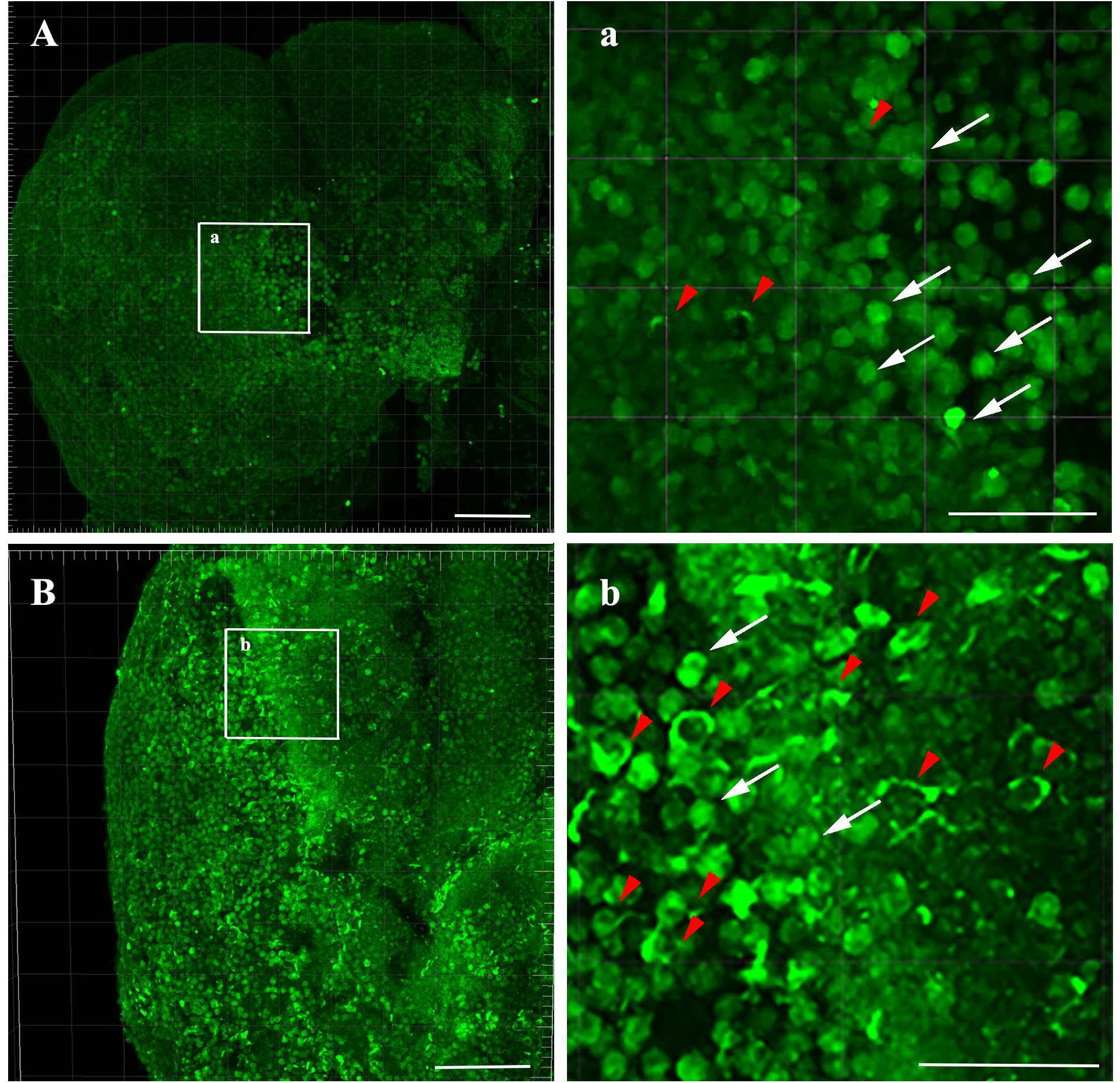
Whole mount staining of TDP-43 in *N. furzeri* brains. Using AbSca/e clarification protocol we were able to observe TDP-43 expression in whole mount brains of young and old *N. furzeri*. In these images, telencephalic regions are shown to clarify how the amount of doughnut-like cells (red arrowheads) is greater in old telencephalon (panel B) as compared to the young animals (panel A). Scale bars indicate 50 μm in each panel.

We then performed a double staining on brain sections, by combining TDP-43 immunofluorescence (**Figure 7, green**) with the Aggresome dye (**Figure 7, red**), to discriminate between generic protein aggregates and specific TDP-43 aggregates, visualized as co-localised green-red fluorescent dots. Comparison of sections from young and old brain tissues showed the presence of several double stained TDP-43 granules in the old tissue (**Figure 7B, b red arrowheads**). Some of the granules were strictly associated to doughnut-like cells. In contrast, consistent with our previous results (Kelmer Sacramento et al., 2020) we were unable to detect aggregates in the young brain tissue, either with or without TDP-43 staining (**Figure 7A**). The red fluorescent signal detectable in the young tissue was not a specific Aggresome staining, but it was to ascribe to auto-fluorescent blood vessels and erythrocyte cells as it is evident in all fluorescent channels.

**Figure 7 –.**
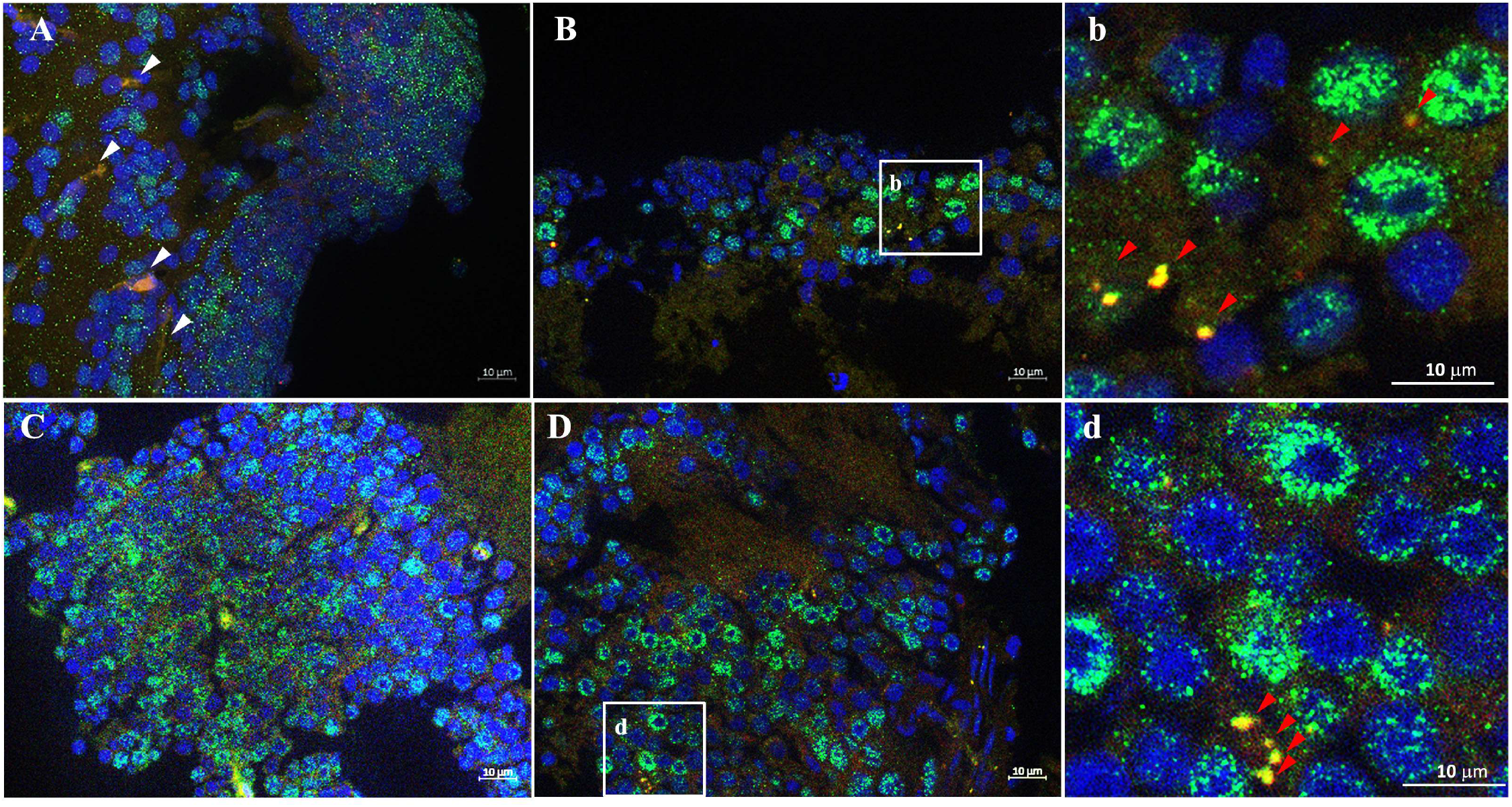
Immunostaining of TDP-43 and colocalization with protein aggregates in old versus young *N. furzeri* brains. Protein aggregates were stained using aggresome dye (red), while TDP-43 was labelled with the monoclonal antibody (green). Both in telencephalon (panels A,B) and optic tectum (panels C,D) we were able to observe aggregates presence in old brains only (panels B,D) while no trace of aggregation was detectable in young samples (panels A,C). Moreover, TDP-43 staining often colocalizes with aggregates signaling, and such aggregates appear to be localized mainly near doughnut-like cells (b/d magnifications, red arrowheads). White arrowheads in panel A indicates blood cells autofluorescence. Scale bars indicate 10 μm in each panel.

These results demonstrate that it is possible to follow ageing-related aggregation of TDP-43 in *N. furzeri* mimicking the neuronal alterations typical of ALS/FTD, and supported the killifish as a convenient model of TDP-43 aggregation in vertebrates with compressed lifespan.

### TDP-43 aggregates localise in stress granules

Finally, we investigated the possible co-localization of TDP-43 with stress granules in *N. furzeri* tissues in agreement with our *in silico* predictions. To do so, we performed double immunostaining for *N. furzeri* TDP-43 and G3BP, a core protein of the stress granules often used as a marker (Martin and Tazi, 2014). The labelling for G3BP was more widespread than that of TDP-43 both in telencephalon and optic tectum, being the signal distributed widely in the cytoplasm of the majority of cells (**Figure 8**). We were nonetheless able to observe the presence of granular structures double-labelled for G3BP and TDP-43 in samples of both young and old animals. This evidence supports the idea that a TDP-43 involvement in stress granules formation and regulation is conserved in *N. furzeri*. Even more importantly, the amount of granules and the dimension of the TDP-43 inclusions seemed to be greater in the samples from old animals **(Figure 8 inset, white arrowheads)**. This is consistent with the idea of an increase of stressing conditions during ageing and thus an increased formation of stress granules inside the cell. Accurate quantification of this visible difference was not attempted for the time being. These exciting results demonstrate the possibility to study granule formation in *N. furzeri*.

**Figure 8 –.**
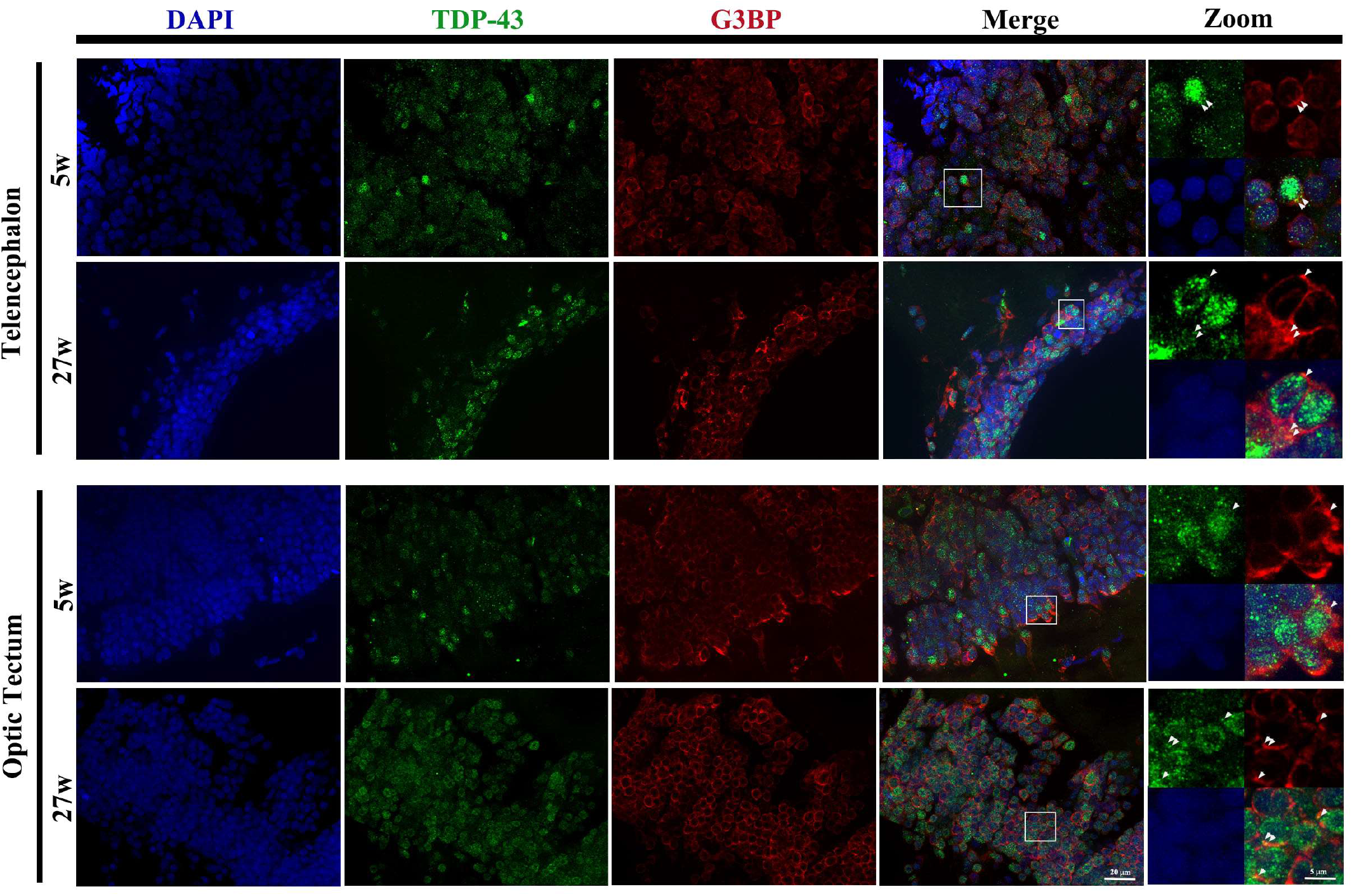
Immunostaining and colocalization of TDP-43 (green) and G3BP (red) in *N. furzeri* tissues. We were able to observe several cells showing partial overlapping signal between TDP-43 and G3BP (white arrowheads in the magnifications of the right column), used here as stress granules marker. Stress granules are present both in tissues of young and old fishes, and are visible in both the optic tectum and telencephalon. A scale bar of 20 μm is the same in all panels of DAPI, TDP-43, G3BP and Merge columns. A scale bar of 5 μm is the same in all panels of the zoom column.

## Discussion

In this manuscript, we laid the foundations for the use of *N. furzeri* as a new animal model to follow the age-dependent aggregation of TDP-43. Increasing evidence has shown that *N. furzeri* shares molecular, histopathological and behavioural ageing-related signatures with mammals making this fish the currently most promising vertebrate model for research on ageing (Cellerino et al., 2016). *N. furzeri* has undoubted advantages as compared to other animal models. It is easily manageable, relatively inexpensive and has a very limited lifespan (3-to-7 months depending on the strain). It is also an animal that presents clear signs of ageing: old fishes appear emaciated, with their spine curved and almost discolored while showing cognitive and locomotor age-dependent decay at a behavioural level. This makes easy to follow the ageing process and relate it to other phenotypes. The possibility to insert human pathogenic mutations in the *N. furzeri* orthologue gene using CRISPR/Cas9 adds up a further advantage in support to this model (Harel et al., 2015).

A bioinformatic analysis demonstrated a remarkably high homology between *N. furzeri* and mammalian protein sequences. Accordingly, we found an amino acidic identity between TDP-43 from *N. furzeri* and Hsa_TDP-43 above 75%. The homology is higher in the N-terminus but breaks down in the C-terminus. This preliminary analysis made us to expect in principle similar properties of nucleocytoplasmic transport, RNA and DNA binding and phase separation as also supported by our *cat*Granule predictions.

We used *in vitro* studies to compare the individual behaviour of different regions of the proteins from human and *N. furzeri* using recombinant fragments spanning the C-termini and the tandem RRM domains. We observed a strong tendency to aggregate of the C-termini in agreement with what has been observed for the human protein (Vega et al., 2019; Capitini et al., 2020). This region is so aggregation prone that the recombinant human C-terminus could be obtained in a soluble form only by cleaving off the glycine-rich region in agreement with previous studies (Capitini et al., 2020). At variance, constructs of the *N. furzeri* C-termini from the end of RRM2 to the end of the proteins are soluble in *E. coli* expression thanks to a shorter glycine-rich region and to a different amino acid composition. This solubility of the *N. furzeri* constructs may constitute an advantage in future studies. Comparison of the behaviour of the *N. furzeri* proteins with Hsa_TDP-43 C-terminus could be achieved by leaving the constructs fused to a SUMO tag or cleaving them in the presence of ThT. We observed a remarkably similar behaviour with the Nfu_TDP-43L C-terminus behaving more similarly to the human protein.

When the analysis was carried out on the RRM1-2 constructs, we predictably noticed only minor differences amongst their aggregation properties and irreversible conformational transitions towards β-rich conformations upon thermal stress. This behaviour demonstrates once again that not only the C-terminus of TDP-43 is involved in protein aggregation and conformational transitions as already observed for the human protein (Zacco et al., 2018). If proteins from organisms as far apart as human and killifish retain a similar tendency to aggregate and misfold, the property must be inherent to the protein and its function.

We then analysed *in silico* the RNA-binding specificities of the full-length proteins. We were prepared to observe almost identical profiles. We found instead only a partial overlap of putative partner sequences, with Nfu_TDP-43L being closer to Hsa-TDP-43. While these results await for a solid experimental confirmation, possibly by iCLIP studies or other cellular screenings, our data could suggest an involvement of regions outside the RRM domains in RNA binding that could explain the presence of more than one TDP-43 in *N. furzeri* and/or different RNA propensities dictated by the minute but appreciable differences in the RRM1-2 sequences of the proteins.

This second hypothesis was directly backed up by our studies. We tested the effects on the aggregation of the *N. furzeri* RRM1-2 domains of the RNA12 aptamer known to bind to Hsa_TDP-43 with affinity in the low nanomolar range (Lukavsky et al., 2013). We have previously demonstrated that the presence of this aptamer is sufficient to increase the protein solubility and inhibit aggregation (Zacco et al., 2019). This has led us to suggest RNA as a cellular chaperone of TDP-43 (Louka et al., 2020). We observed also for the *N. furzeri* proteins an appreciable reduction of aggregation. Interestingly, the effect is less marked for Hsa_TDP-43 and Nfu_TDP-43L than for Nfu_TDP-43, suggesting a lower affinity, either due to a different mode of interaction or, more likely, to a less tight surface of interaction. Given the high degree of homology and the identity of almost all residues in contact with the aptamer as observed in the solution structure of the complex, we suggest that a lower affinity to the aptamer could be caused by the six amino acid insertion between the two RRM domains of Nfu_TDP-43 (**Figure 1**). A different length of the linker could change the mutual orientation of the two domains and their dynamics which could in turn affect the interaction with RNA. It is interesting to note that the insertion is immediately after R181, the residue associated with a recently described clinically important mutation (Chen et al., 2019). Together, these observations point at a previously unreported important role in RNA recognition of the linker between the two RRM domains.

In parallel to these *in silico* and *in vitro* studies, we carried out an *ex vivo* investigation in animal tissues to test whether TDP-43 can form aggregates also in the animal and/or behaves differently in young and old fishes. Using immunofluorescence experiments, we proved that *N. furzeri* cells may show an abnormal distribution of nuclear TDP-43 (doughnut-like cells) that increases markedly during ageing. This could be interpreted as a sign of impaired nucleo-cytoplasmatic transport, which does not come as a surprise since dysfunction of nucleo-cytoplasmatic transport is often associated to neurodegenerative diseases and is also observed in physiological ageing (Hutten and Dormann, 2020). TDP-43 neurotoxicity in neurodegenerative disorders could thus be linked to a loss of function of nuclear TDP-43 (Xu, 2012) and, more in general, to dysfunction of nucleo-cytoplasmatic transport. We were then able to observe protein aggregates exclusively in old brain tissues. Some of these aggregates were positive to TDP-43 staining and strictly associated to doughnut-like cells. This behaviour allowed us to conclude that *N. furzeri* TDP-43 is indeed able to form intracellular pathological aggregates *in vivo*, and strengthened the interpretation of doughnut-like cells as a pathological condition that can, in the future, be used as a marker to identify aged animals.

We also demonstrated co-localization of TDP-43 and G3BP, both in young and old animals, confirming that the interaction between TDP-43 and stress granules is a common feature also in *N. furzeri*. It has been shown that stress granules impair nucleocytoplasmatic transport and that suppression of stress granules prevents neurodegeneration in a ALS/FTD model based on C9ORF72 (Zhang et al., 2018). This observation suggests that, by preventing access of TDP-43 to the nucleus, stress granule formation may be central to the pathogenic mechanisms of ALS. It will thus be of great interest to study such interactions further in this animal model and find ways to link stress granule dynamics dysregulation with TDP-43 related neurodegenerative diseases.

In conclusion, our results clearly demonstrate the feasibility of using *N. furzeri* as a powerful model for TDP-43-related diseases, exploiting the advantages of the short lifespan of this organism, within the limitations intrinsic in any model. The small but interesting differences in the aggregation and RNA-binding properties of the individual sequences may also be favourably exploited in the future to deepen our understanding of protein aggregation, stress granule formation, RNA specificity and neurodegenerative processes. Further studies will be needed to observe the TDP-43-related ageing process in *N. furzeri* in a more statistically significant manner and to identify possible modifiers of the process.

## Acknowledgements

This work was supported by the Dementia Research UK (RE1 3556) and by the Francis Crick Institute through provision of access to the MRC Biomedical NMR Centre. The Francis Crick Institute receives its core funding from Cancer Research UK (FC001029), the UK Medical Research Council (FC001029), and the Wellcome Trust (FC001029). GGT’s research was supported by European Research Council (RIBOMYLOME_309545 and ASTRA_855923), the H2020 projects IASIS_727658 and INFORE_825080, the Spanish Ministry of Economy and Competitiveness BFU2017-86970-P, the European Union’s Horizon 2020 research and innovation programme under the Marie Skłodowska-Curie grant agreement No 754490 within the MINDED project.

## References

Agostini, F., Cirillo, D., Ponti, R. D., and Tartaglia, G. G. (2014). SeAMotE: a method for high-throughput motif discovery in nucleic acid sequences. BMC Genomics 15, 925. doi:10.1186/1471-2164-15-925.

Agostini, F., Zanzoni, A., Klus, P., Marchese, D., Cirillo, D., and Tartaglia, G. G. (2013). catRAPID omics: a web server for large-scale prediction of protein-RNA interactions. Bioinformatics 29, 2928–2930. doi:10.1093/bioinformatics/btt495.

Alami, N. H., Smith, R. B., Carrasco, M. A., Williams, L. A., Winborn, C. S., Han, S. S. W., et al. (2014). Axonal Transport of TDP-43 mRNA Granules Is Impaired by ALS-Causing Mutations. Neuron 81, 536–543. doi:10.1016/j.neuron.2013.12.018.

Anderson, P., and Kedersha, N. (2009). RNA granules: post-transcriptional and epigenetic modulators of gene expression. Nat. Rev. Mol. Cell Biol. 10, 430–436. doi:10.1038/nrm2694.

Ayala, Y. M., De Conti, L., Avendaño-Vázquez, S. E., Dhir, A., Romano, M., D’Ambrogio, A., et al. (2011). TDP-43 regulates its mRNA levels through a negative feedback loop. EMBO J. 30, 277–288. doi:10.1038/emboj.2010.310.

Baumgart, M., Priebe, S., Groth, M., Hartmann, N., Menzel, U., Pandolfini, L., et al. (2016). Longitudinal RNA-Seq Analysis of Vertebrate Aging Identifies Mitochondrial Complex I as a Small-Molecule-Sensitive Modifier of Lifespan. Cell Syst. 2, 122–132. doi:10.1016/j.cels.2016.01.014.

Bellucci, M., Agostini, F., Masin, M., and Tartaglia, G. G. (2011). Predicting protein associations with long noncoding RNAs. Nat. Methods 8, 444–445. doi:10.1038/nmeth.1611.

Berning, B. A., and Walker, A. K. (2019). The Pathobiology of TDP-43 C-Terminal Fragments in ALS and FTLD. Front. Neurosci. 13, 335. doi:10.3389/fnins.2019.00335.

Bhardwaj, A., Myers, M. P., Buratti, E., and Baralle, F. E. (2013). Characterizing TDP-43 interaction with its RNA targets. Nucleic Acids Res. 41, 5062–5074. doi:10.1093/nar/gkt189.

Bolognesi, B., Lorenzo Gotor, N., Dhar, R., Cirillo, D., Baldrighi, M., Tartaglia, G. G., et al. (2016). A Concentration-Dependent Liquid Phase Separation Can Cause Toxicity upon Increased Protein Expression. Cell Rep. 16, 222–231. doi:10.1016/j.celrep.2016.05.076.

Buratti, E., and Baralle, F. E. (2001). Characterization and functional implications of the RNA binding properties of nuclear factor TDP-43, a novel splicing regulator of CFTR exon 9. J. Biol. Chem. 276, 36337–36343. doi:10.1074/jbc.M104236200.

Capitini, C., Fani, G., Vivoli Vega, M., Penco, A., Canale, C., Cabrita, L. D., et al. (2020). Full-length TDP-43 and its C-terminal domain form filaments in vitro having nonamyloid properties. Amyloid, 1–10. doi:10.1080/13506129.2020.1826425.

Cellerino, A., Valenzano, D. R., and Reichard, M. (2016). From the bush to the bench: the annual Nothobranchius fishes as a new model system in biology. Biol. Rev. 91, 511–533. doi:10.1111/brv.12183.

Chen, H.-J., Topp, S. D., Hui, H. S., Zacco, E., Katarya, M., McLoughlin, C., et al. (2019). RRM adjacent TARDBP mutations disrupt RNA binding and enhance TDP-43 proteinopathy. Brain 142, 3753–3770. doi:10.1093/brain/awz313.

Chu, J.-F., Majumder, P., Chatterjee, B., Huang, S.-L., and Shen, C.-K. J. (2019). TDP-43 Regulates Coupled Dendritic mRNA Transport-Translation Processes in Co-operation with FMRP and Staufen1. Cell Rep. 29, 3118–3133.e6. doi:10.1016/j.celrep.2019.10.061.

Cui, R., Willemsen, D., and Valenzano, D. R. (2020). <em>Nothobranchius *N. furzeri* (African Turquoise Killifish). Trends Genet. 36, 540–541. doi:10.1016/j.tig.2020.01.012.

Dolfi, L., Ripa, R., Antebi, A., Valenzano, D. R., and Cellerino, A. (2019). Cell cycle dynamics during diapause entry and exit in an annual killifish revealed by FUCCI technology. Evodevo 10, 29. doi:10.1186/s13227-019-0142-5.

Gibson, D. G., Young, L., Chuang, R.-Y., Venter, J. C., Hutchison, C. A. 3rd, and Smith, H. O. (2009). Enzymatic assembly of DNA molecules up to several hundred kilobases. Nat. Methods 6, 343–345. doi:10.1038/nmeth.1318.

Gilks, N., Kedersha, N., Ayodele, M., Shen, L., Stoecklin, G., Dember, L. M., et al. (2004). Stress granule assembly is mediated by prion-like aggregation of TIA-1. Mol. Biol. Cell 15, 5383–5398. doi:10.1091/mbc.e04-08-0715.

Gotor, N. L., Armaos, A., Calloni, G., Vabulas, R. M., de Groot, N. S., and Tartaglia, G. G. (2020). RNA-Binding and Prion Domains: The Yin and Yang of Phase Separation. Nucleic Acids Res., 10.1093/nar/gkaa681. doi:10.1101/2020.01.14.904383.

Hama, H., Hioki, H., Namiki, K., Hoshida, T., Kurokawa, H., Ishidate, F., et al. (2015). ScaleS: an optical clearing palette for biological imaging. Nat. Neurosci. 18, 1518–1529. doi:10.1038/nn.4107.

Harel, I., Benayoun, B. A., Machado, B., Singh, P. P., Hu, C.-K., Pech, M. F., et al. (2015). A Platform for Rapid Exploration of Aging and Diseases in a Naturally Short-Lived Vertebrate. Cell 160, 1013–1026. doi:10.1016/j.cell.2015.01.038.

Hergesheimer, R. C., Chami, A. A., de Assis, D. R., Vourc’h, P., Andres, C. R., Corcia, P., et al. (2019). The debated toxic role of aggregated TDP-43 in amyotrophic lateral sclerosis: a resolution in sight? Brain 142, 1176–1194. doi:10.1093/brain/awz078.

Hu, C.-K., Wang, W., Brind’Amour, J., Singh, P. P., Reeves, G. A., Lorincz, M. C., et al. (2020). Vertebrate diapause preserves organisms long term through Polycomb complex members. Science (80-.). 367, 870 LP–874. doi:10.1126/science.aaw2601.

Hutten, S., and Dormann, D. (2020). Nucleocytoplasmic transport defects in neurodegeneration - Cause or consequence? Semin. Cell Dev. Biol. 99, 151–162. doi:10.1016/j.semcdb.2019.05.020.

Ishiguro, A., Kimura, N., Watanabe, Y., Watanabe, S., and Ishihama, A. (2016). TDP-43 binds and transports G-quadruplex-containing mRNAs into neurites for local translation. Genes to Cells 21, 466–481. doi:10.1111/gtc.12352.

Kelmer Sacramento, E., Kirkpatrick, J. M., Mazzetto, M., Baumgart, M., Bartolome, A., Di Sanzo, S., et al. (2020). Reduced proteasome activity in the aging brain results in ribosome stoichiometry loss and aggregation. Mol. Syst. Biol. 16, e9596. doi:10.15252/msb.20209596.

Kim, S. H., Shanware, N. P., Bowler, M. J., and Tibbetts, R. S. (2010). Amyotrophic lateral sclerosis-associated proteins TDP-43 and FUS/TLS function in a common biochemical complex to co-regulate HDAC6 mRNA. J. Biol. Chem. 285, 34097–34105. doi:10.1074/jbc.M110.154831.

Lang, B., Armaos, A., and Tartaglia, G. G. (2019). RNAct: Protein-RNA interaction predictions for model organisms with supporting experimental data. Nucleic Acids Res. 47, D601–D606. doi:10.1093/nar/gky967.

Li, H.-R., Chen, T.-C., Hsiao, C.-L., Shi, L., Chou, C.-Y., and Huang, J. (2018a). The physical forces mediating self-association and phase-separation in the C-terminal domain of TDP-43. Biochim. Biophys. Acta - Proteins Proteomics 1866, 214–223. doi:https://doi.org/10.1016/j.bbapap.2017.10.001.

Li, H.-R., Chiang, W.-C., Chou, P.-C., Wang, W.-J., and Huang, J.-R. (2018b). TAR DNA-binding protein 43 (TDP-43) liquid-liquid phase separation is mediated by just a few aromatic residues. J. Biol. Chem. 293, 6090–6098. doi:10.1074/jbc.AC117.001037.

Liu-Yesucevitz, L., Bilgutay, A., Zhang, Y.-J., Vanderwyde, T., Citro, A., Mehta, T., et al. (2010). Tar DNA Binding Protein-43 (TDP-43) Associates with Stress Granules: Analysis of Cultured Cells and Pathological Brain Tissue. PLoS One 5, e13250. Available at: https://doi.org/10.1371/journal.pone.0013250.

Liu, Y.-C., Chiang, P.-M., and Tsai, K.-J. (2013). Disease animal models of TDP-43 proteinopathy and their pre-clinical applications. Int. J. Mol. Sci. 14, 20079–20111. doi:10.3390/ijms141020079.

Livi, C. M., Klus, P., Delli Ponti, R., and Tartaglia, G. G. (2016). catRAPID signature: identification of ribonucleoproteins and RNA-binding regions. Bioinformatics 32, 773–775. doi:10.1093/bioinformatics/btv629.

Loganathan, S., Lehmkuhl, E. M., Eck, R. J., and Zarnescu, D. C. (2020). To Be or Not To Be.. .Toxic—Is RNA Association With TDP-43 Complexes Deleterious or Protective in Neurodegeneration?. Front. Mol. Biosci. 6, 154. Available at: https://www.frontiersin.org/article/10.3389/fmolb.2019.00154.

Louka, A., Zacco, E., Temussi, P. A., Tartaglia, G. G., and Pastore, A. (2020). RNA as the stone guest of protein aggregation. Nucleic Acids Res. 48, 11880–11889. doi:10.1093/nar/gkaa822.

Lukavsky, P. J., Daujotyte, D., Tollervey, J. R., Ule, J., Stuani, C., Buratti, E., et al. (2013). Molecular basis of UG-rich RNA recognition by the human splicing factor TDP-43. Nat. Struct. Mol. Biol. 20, 1443–1449. doi:10.1038/nsmb.2698.

Mackenzie, I. R. A., and Rademakers, R. (2008). The role of transactive response DNA-binding protein-43 in amyotrophic lateral sclerosis and frontotemporal dementia. Curr. Opin. Neurol. 21, 693–700. doi:10.1097/WCO.0b013e3283168d1d.

Martin, S., and Tazi, J. (2014). Visualization of G3BP stress granules dynamics in live primary cells. J. Vis. Exp., 51197. doi:10.3791/51197.

Matsui, H., Kenmochi, N., and Namikawa, K. (2019). Age-and α-Synuclein-Dependent Degeneration of Dopamine and Noradrenaline Neurons in the Annual Killifish Nothobranchius *N. furzeri*. Cell Rep. 26, 1727–1733.e6. doi:https://doi.org/10.1016/j.celrep.2019.01.015.

McDonald, K. K., Aulas, A., Destroismaisons, L., Pickles, S., Beleac, E., Camu, W., et al. (2011). TAR DNA-binding protein 43 (TDP-43) regulates stress granule dynamics via differential regulation of G3BP and TIA-1. Hum. Mol. Genet. 20, 1400–1410. doi:10.1093/hmg/ddr021.

Mompeán, M., Buratti, E., Guarnaccia, C., Brito, R. M. M., Chakrabartty, A., Baralle, F. E., et al. (2014). “Structural characterization of the minimal segment of TDP-43 competent for aggregation”. Arch. Biochem. Biophys. 545, 53–62. doi:10.1016/j.abb.2014.01.007.

Mompeán, M., Hervás, R., Xu, Y., Tran, T. H., Guarnaccia, C., Buratti, E., et al. (2015). Structural Evidence of Amyloid Fibril Formation in the Putative Aggregation Domain of TDP-43. J. Phys. Chem. Lett. 6, 2608–2615. doi:10.1021/acs.jpclett.5b00918.

Neelagandan, N., Gonnella, G., Dang, S., Janiesch, P. C., Miller, K. K., Küchler, K., et al. (2018). TDP-43 enhances translation of specific mRNAs linked to neurodegenerative disease. Nucleic Acids Res. 47, 341–361. doi:10.1093/nar/gky972.

Neumann, M., Sampathu, D. M., Kwong, L. K., Truax, A. C., Micsenyi, M. C., Chou, T. T., et al. (2006). Ubiquitinated TDP-43 in frontotemporal lobar degeneration and amyotrophic lateral sclerosis. Science 314, 130–133. doi:10.1126/science.1134108.

Ou, S. H., Wu, F., Harrich, D., García-Martínez, L. F., and Gaynor, R. B. (1995). Cloning and characterization of a novel cellular protein, TDP-43, that binds to human immunodeficiency virus type 1 TAR DNA sequence motifs. J. Virol. 69, 3584–3596.

Parker, S. J., Meyerowitz, J., James, J. L., Liddell, J. R., Crouch, P. J., Kanninen, K. M., et al. (2012). Endogenous TDP-43 localized to stress granules can subsequently form protein aggregates. Neurochem. Int. 60, 415–424. doi:10.1016/j.neuint.2012.01.019.

Philippe, C., Hautekiet, P., Grégoir, A. F., Thoré, E. S. J., Pinceel, T., Stoks, R., et al. (2018). Combined effects of cadmium exposure and temperature on the annual killifish (Nothobranchius *N. furzeri)*. Environ. Toxicol. Chem. 37, 2361–2371. doi:10.1002/etc.4182.

Polymenidou, M., Lagier-Tourenne, C., Hutt, K. R., Huelga, S. C., Moran, J., Liang, T. Y., et al. (2011). Long pre-mRNA depletion and RNA missplicing contribute to neuronal vulnerability from loss of TDP-43. Nat. Neurosci. 14, 459–468. doi:10.1038/nn.2779.

Prasad, A., Bharathi, V., Sivalingam, V., Girdhar, A., and Patel, B. K. (2019). Molecular Mechanisms of TDP-43 Misfolding and Pathology in Amyotrophic Lateral Sclerosis. Front. Mol. Neurosci. 12, 25. doi:10.3389/fnmol.2019.00025.

Reichwald, K., Petzold, A., Koch, P., Downie, B. R., Hartmann, N., Pietsch, S., et al. (2015). Insights into Sex Chromosome Evolution and Aging from the Genome of a Short-Lived Fish. Cell 163, 1527–1538. doi:10.1016/j.cell.2015.10.071.

Sephton, C. F., Good, S. K., Atkin, S., Dewey, C. M., Mayer 3rd, P., Herz, J., et al. (2010). TDP-43 is a developmentally regulated protein essential for early embryonic development. J. Biol. Chem. 285, 6826–6834. doi:10.1074/jbc.M109.061846.

Shen, D., Coleman, J., Chan, E., Nicholson, T. P., Dai, L., Sheppard, P. W., et al. (2011). Novel Cell-and Tissue-Based Assays for Detecting Misfolded and Aggregated Protein Accumulation Within Aggresomes and Inclusion Bodies. Cell Biochem. Biophys. 60, 173–185. doi:10.1007/s12013-010-9138-4.

Shih, Y.-H., Tu, L.-H., Chang, T.-Y., Ganesan, K., Chang, W.-W., Chang, P.-S., et al. (2020). TDP-43 interacts with amyloid-β, inhibits fibrillization, and worsens pathology in a model of Alzheimer’s disease. Nat. Commun. 11, 5950. doi:10.1038/s41467-020-19786-7.

Terzibasi, E., Valenzano, D. R., Benedetti, M., Roncaglia, P., Cattaneo, A., Domenici, L., et al. (2008). Large Differences in Aging Phenotype between Strains of the Short-Lived Annual Fish Nothobranchius *N. furzeri*. PLoS One 3, e3866. Available at: https://doi.org/10.1371/journal.pone.0003866.

Tollervey, J. R., Curk, T., Rogelj, B., Briese, M., Cereda, M., Kayikci, M., et al. (2011). Characterizing the RNA targets and position-dependent splicing regulation by TDP-43. Nat. Neurosci. 14, 452–458. doi:10.1038/nn.2778.

Tourrière, H., Chebli, K., Zekri, L., Courselaud, B., Blanchard, J. M., Bertrand, E., et al. (2003). The RasGAP-associated endoribonuclease G3BP assembles stress granules. J. Cell Biol. 160, 823–831. doi:10.1083/jcb.200212128.

Tozzini, E. T., Baumgart, M., Battistoni, G., and Cellerino, A. (2012). Adult neurogenesis in the short-lived teleost Nothobranchius *N. furzeri:* localization of neurogenic niches, molecular characterization and effects of aging. Aging Cell 11, 241–251. doi:10.1111/j.1474-9726.2011.00781.x.

Valdesalici, S., and Cellerino, A. (2003). Extremely short lifespan in the annual fish Nothobranchius *N. furzeri*. Proc. R. Soc. London. Ser. B Biol. Sci. 270, S189–S191. doi:10.1098/rsbl.2003.0048.

Valenzano, D. R., Benayoun, B. A., Singh, P. P., Zhang, E., Etter, P. D., Hu, C.-K., et al. (2015). The African Turquoise Killifish Genome Provides Insights into Evolution and Genetic Architecture of Lifespan. Cell 163, 1539–1554. doi:10.1016/j.cell.2015.11.008.

Valenzano, D. R., Terzibasi, E., Genade, T., Cattaneo, A., Domenici, L., and Cellerino, A. (2006). Resveratrol prolongs lifespan and retards the onset of age-related markers in a short-lived vertebrate. Curr. Biol. 16, 296–300. doi:10.1016/j.cub.2005.12.038.

Vega, M. V., Nigro, A., Luti, S., Capitini, C., Fani, G., Gonnelli, L., et al. (2019). Isolation and characterization of soluble human full-length TDP-43 associated with neurodegeneration. FASEB J. 33, 10780–10793. doi:https://doi.org/10.1096/fj.201900474R.

Vrtílek, M., Žák, J., Pšenička, M., and Reichard, M. (2018). Extremely rapid maturation of a wild African annual fish. Curr. Biol. 28, R822–R824. doi:10.1016/j.cub.2018.06.031.

Wendler, S., Hartmann, N., Hoppe, B., and Englert, C. (2015). Age-dependent decline in fin regenerative capacity in the short-lived fish Nothobranchius *N. furzeri*. Aging Cell 14, 857–866. doi:10.1111/acel.12367.

Xu, Z.-S. (2012). Does a loss of TDP-43 function cause neurodegeneration? Mol. Neurodegener. 7, 27. doi:10.1186/1750-1326-7-27.

Zacco, E., Graña-Montes, R., Martin, S. R., de Groot, N. S., Alfano, C., Tartaglia, G. G., et al. (2019). RNA as a key factor in driving or preventing self-assembly of the TAR DNA-binding protein 43. J. Mol. Biol. 431, 1671–1688. doi:https://doi.org/10.1016/j.jmb.2019.01.028.

Zacco, E., Martin, S. R., Thorogate, R., and Pastore, A. (2018). The RNA-Recognition Motifs of TAR DNA-Binding Protein 43 May Play a Role in the Aberrant Self-Assembly of the Protein. Front. Mol. Neurosci. 11, 372. Available at: https://www.frontiersin.org/article/10.3389/fnmol.2018.00372.

Zhang, K., Daigle, J. G., Cunningham, K. M., Coyne, A. N., Ruan, K., Grima, J. C., et al. (2018). Stress Granule Assembly Disrupts Nucleocytoplasmic Transport. Cell 173, 958–971.e17. doi:10.1016/j.cell.2018.03.025.

